# Traditional uses, population, threats and conservation of the bansouman or gingerbread plum *Neocarya macrophylla* (Chrysobalanaceae) in Republic of Guinea (West Africa)

**DOI:** 10.1101/2025.06.08.655547

**Authors:** Denise Molmou, Martin Cheek, Gbamon Konomou, Sékou Magassouba, Ana Rita Simoes, Isabel Larridon, Charlotte Couch

## Abstract

The flora of the Republic of Guinea (West Africa) includes nearly 4000 vascular plants species, of which c. 10% have been documented to have local socio-economic uses. These socio-economic plants provide among others, foods, medicines, materials and a source of income for the people of Guinea. One example is the bansouman or gingerbread plum, *Neocarya macrophylla* (Chrysobalanaceae) which is locally valued for its large, delicious seeds (’nuts’), edible fruit flesh, and medicinal bark (among other medicinal parts). Through a field survey of 17 local communities adjacent to *Neocarya* populations and the *Neocarya* populations themselves (780 mature individuals measured), in seven of the ten Guinean prefectures from which the species is recorded, this study documents (1) the local names, harvesting methods, uses and seed sale prices at different points in the value chain, partitions the species by habitat, records perceived threats to trees, key elements of the population structure and relative frequency of *Neocarya macrophylla* in Guinea, (2) map and re assesses its global conservation status as Near Threatened using the 2012 IUCN Red List categories and criteria, and (3) construct a species conservation action plan.

Analysis of our survey data suggests that the largest number of mature trees (75% of the total) is in the three northern prefectures (Gaoual, Koundara, Pita) at the western foot of the Fouta massif whence seeds are exported internationally. Trees mainly occurred in shrub savannah (45% of mature individuals), with lower numbers in three other habitats. Human set bush fires appear to be the most ubiquitous threat (55%) in Guinea, but sand quarrying (18%), agricultural conversion and debarking are also key threats. Natural regeneration levels seem low compared with data from Niger with juveniles (<1 m tall) less than 10 % of the number of mature, fruiting trees (> 11 cm circumference) recorded.

This is the first study focused on *Neocarya macrophylla* in the Republic of Guinea. In contrast the species has been the subject of numerous other papers elsewhere in its range, and we document 14 studies from Niger alone in the last 15 years. While the species is well known as an important Indigenous Fruit Tree with multiple uses in some countries, it is currently not among the top 10 such species listed for Guinea.

Compared with Nigeria where numerous other uses including industrial are derived from the species, *Neocarya macrophylla* appears under utilised in Guinea.

We suggest that studies of the chemistry and nutritional value of the seeds of Guinean *Neocarya* are made, since there are indications that these differ between Guinea and e.g. Niger. Follow up studies to map tree distribution in Guinea in more detail, quantify tree density, fruit yield per tree and per Ha, and duration between germination and fruiting are advised. Field trials to select elite individuals are recommended. Development of a machine to extract seeds from the endocarp (decortication) is a priority as this is the major factor limiting wild harvesting and availability of seed for human consumption.

## Introduction

The flora of the Republic of Guinea (West Africa) comprises nearly 4000 species (including subspecies) of vascular plants (Gosline *et al*. 2023a; 2023b). Among these, 399 indigenous species are considered to have socio-economic uses (Molmou *et al*. submitted). Useful plant species provide an important contribution globally to local economies in terms of value and resources (Shelef *et al*. 2017). Native tree species are an integral part of production systems in many village communities (Jepsen *et al*. 2024). Some farmers in West Africa preserve wild tree species such as the gingerbread plum (*Neocarya macrophylla* (Sabine) Prance ex F.White), niri (*Parkia bicolor* A.Chev.) and the mana or shea tree (*Vitellaria paradoxa* C.F.Gaertn.), in their agricultural fields for their socio-economic uses (Guimbo *et al*. 2017; Bayala & Harmand 2023). Indeed, the harvesting and processing of products from spontaneously occurring native plant species provides a real opportunity for rural communities, especially women and children (Guimbo *et al*. 2017).

More than 250 threatened plant species have been documented in Guinea (https://www.iucnredlist.org/, accessed Oct. 2024; Couch *et al*. (2019a)). Their protection should not only be a national responsibility, but a global priority. Along with this significant floral wealth, resources such as bauxite, high-grade iron ore reserves, as well as diamonds, gold, but also granite and sand, leads to risks from open-cast mining and quarrying and so pressure on species habitats increases with the need to generate revenue in the country (Couch *et al*. 2023). This is in addition to unsustainable slash-and-burn agriculture, logging, and population growth, which rely heavily on firewood and charcoal for fuel (Couch *et al*. 2019b). Fires set to rejuvenate grasslands for grazing and clear areas for agriculture pose a serious threat to threatened habitats in Guinea, including forests (Cheek *et al*. 2020). Indeed, each year the country burns from top to bottom. Wildfires are a growing problem, and smoke significantly affects air quality (Garrett-Bakelman *et al*. 2019). However, the goal of conservationists is to protect globally threatened species and achieve a successful recovery of them, documenting them on the green list (Akçakaya *et al*. 2018).

*Neocarya macrophylla*, formerly known *as Parinari macrophylla* Sabine, belongs to the Chrysobalanaceae family. For a full description of the species see Prance & Sothers (2003). The genus with its single species has been shown by molecular phylogenetic studies to be sister to pantropical *Parinari* Aubl. (Bardon *et al*. 2013; Chave *et al*. 2020). It is listed as a socio-economic plant of Guinea (Molmou *et al*. submitted). It is a species of tree that is found from Senegal to northern Nigeria. Records from Cameroon, Sudan and Central African Republic (e.g. POWO continuously updated; gbif.org) appear to be based on misidentifications, e.g. the species does not appear for Sudan in the Checklist of Darbyshire *et al*. 2015). Within its natural range the species occurs mainly on sandy soils along the coast from Senegal to Liberia and, separately, in the northern half of the Sudanian region, 700-1000 km inland (Prance & Sothers 2003). The species has large seeds up to 32 x 13 × 10 mm. These have white flesh with a brown testa*. Neocarya macrophylla* seeds are harvested from the wild and are locally marketed in Guinea, seeds are edible (Burkill, 1985) and crushed to produce cooking oil and edible paste. Phytochemistry studies (e.g. Amza *et al*. 2010, 2011, Garba *et al*. 2022, Imam *et al*. 2023, Bala *et al*. 2022) confirm the presence of diverse seed oils at high levels, proteins at c. 20% and essential and non-essential amino acids and vitamins (B1, B2, B6, and biotin H) in gingerbread plum kernel flour and paste in Niger and Guinea (Yusuf *et al*. 2024). According to Diaby *et al*. (2016), there is significant variation in the seed oil composition and content between trees of Niger and Guinea.

*Neocarya macrophylla* has been studied as an Indigenous Fruit Tree, mainly for its uses, in most countries within its natural range such as Senegal (Sambou *et al*. 2024; Djihounouck *et al*. 2021), Ghana, Nigeria, but it has been most highly studied in S.W. Niger, where 14 studies have been published in the last 15 years: Abdoulahi *et al*. (2022), Aboubacar *et al*. (2018), Balla *et al*. (2008), Diaby *et al* 2016, Garba *et al* (2022), Guimbo *et al*. (2010, 2011, 2016a, 2016b, 2017), Ide *et al*. (2024), Kolafane *et al*. (2018a, 2018b), Naturelles & Maradi. (2017). These cover multiple aspects of the species including the physico chemical, the nutritional, the medicinal, to the structure of populations, the level of natural regeneration and also *in vitro* multiplication. In contrast we have found no papers at all on the uses of the species from Liberia, Sierra Leone, Cote D’Ivoire, Benin, nor Guinea.

*Neocarya macrophylla* is traditionally used as a food and for medicinal, spiritual and industrial purposes in various parts of its range. It is also used as a soap, dye, glue, fodder, termite repellent, firewood and for structural materials in Nigeria (Yusuf *et al*.2024). Recent studies in Niger have reported the importance of the tree for various food and medicinal uses and that the subpopulations of this tree are not in decline in its natural environment, where it can occur at 25 mature trees (and 1118 seedlings) per ha, each tree yielding as much as 390 kg of fruit p.a. (Guimbo *et al*. 2017). In Guinea, some localities where this species previously occurred have disappeared due to clearing for plantations of exotic species such as cashew (*Anacardium occidentale* L.; Molmou *et al*. 2022) supported by a government intervention. A previously unpublished 2023 study of the use of *Neocarya* in Guinea by United Purpose was commissioned by authors of this paper (Appendix 1). This looked at the value chain and utility of *Neocarya* in the prefectures of Koundara, and to a lesser extent of Gaoual. The study showed that fruits are collected mainly by women and their children, from under the trees near their villages after the flesh has been eaten by animals, as well as collected fresh. The main season for fruit collection is January-March inclusive. The fruit and seeds are important in the local diet, and bark and leaves used medicinally. Extraction of seeds from the tough endocarp is time consuming. Sales to collectors for market traders in the towns provides important income to rural women yet fruit can remain on the tree, uncollected, due to low prices. Large but unrecorded quantities of seeds are exported to the capital Conakry, and internationally to Guinea-Bissau, Senegal and Nigeria (United Purpose 2023 Appendix 1).

Here we conducted a study on the traditional use and conservation of *Neocarya macrophylla* in Guinea. This study aims to (1) document the use of *Neocarya macrophylla* in Guinea through a field survey and interviews, (2) assess its global conservation status using the IUCN Red List categories and criteria to understand if its use is sustainable or negatively impacts its conservation status, and (3) compile a species conservation action plan to inform the sustainable use and conservation of the species in Guinea.

## Material and Methods

### Field survey and interviews

Based on a desktop survey using Google Earth and existing occurrence records (e.g. GBIF.Org 2025) we determined in which prefectures the sites are located, targeted suitable sites where the species would likely occur and studied access. A questionnaire on *Neocarya macrophylla* concerning traditional uses, vernacular name, local distribution, harvesting method, sale, and threats to the species in Guinea was prepared (Appendix 2A).

The field survey was conducted in January 2020 with the help of local elected officials and comprised of a) interviews of local communities interacting with wild (sub)populations of *Neocarya* and b) a survey of wild (sub)populations of the species. Interviews were conducted in focus groups of three to five people per village, and participants were selected based on their availability and knowledge of *Neocarya macrophylla*. A total of 84 people were interviewed, including 45 women and 40 men, aged 30 to 55; their occupations were all related to agriculture or work on local land. The study area included 18 villages corresponding to 2–3 sub-prefectures of seven prefectures (see Table 1). One questionnaire was completed for the focus group of each of the 18 villages and the results summarized for each of the seven prefectures surveyed (Appendix 2B).The prefectures of the study area likely contains the majority of the individuals of *Neocarya* in Guinea, however the species is also recorded from the prefectures of Forecariah (*Cheek* 15910, HNG, K) and Boké (*Fofana* 4, HNG, K) although numbers of individuals at these sites are thought to be relatively low. Finally, the species is also known to occur in Koundara prefecture, where numbers of individuals are high (United Purpose 2023, Appendix 1). It was intended to include Koundara in the present survey but logistic difficulties rendered access impossible at the time.

**Table 1.** Villages/sites sampled, with local names of *Neocarya*, number of mature and juvenile plants (trees <11 cm circumf. and juveniles >1 m high were not recorded) and voucher specimens.

| Prefect. | Site name | Local name | No. of mature trees (>11 cm circumf.) observed | No. of juvenile individuals Observed (<1 m tall) | No. of herbarium voucher specimens |
| --- | --- | --- | --- | --- | --- |
| Telimélé | Garama | Bansouman | 4 | 0 | 1 |
|  | Mahbe | Bansouman | 100 | 18 | 30 |
|  | Sinta - Siraya | Bansouman | 70 | 6 | 31 |
|  | Gougouse | Bansouman | 3 | 0 | 1 |
|  | Santou | Bansouman | 20 | 2 | 17 |
|  | Donzon | Bansouman | 11 | 1 | 7 |
| Gaoual | Koumbia N'djouria | Koura-Bansouman | 32 | 2 | 30 |
|  | Koumbia Kithiare | Koura-Bansouman | 100 | 12 | 30 |
|  | Koumbia Wanta | Koura-Bansouman | 5 | 0 | 2 |
| Pita | Dongholtou ma | Bansouman | 8 | 0 | 5 |
|  | Ley Miro | Bansouman | 50 | 5 | 29 |
|  | Faro | Bansouman | 150 | 11 | 31 |
|  | Sangaria | Bansouman | 92 | 4 | 30 |
| Kindia | Mayon Kouré | Gnamouie or Bansouman | 50 | 0 | 7 |
| Boffa | Lefoureboui | Bansouman | 60 | 2 | 30 |
| Dubreka | Tanènè | Bansouman | 15 | 0 | 9 |
| Coyah | Kouria | No name given | 10 | 0 | 1 |
|  | **Total** |  | **780** | **63** | **291** |

The first author obtained permission from village authorities for interviews and individuals were interviewed after seeing the “Ordre de Mission” (Governmental fieldwork permit), signed by regional and prefectoral administrative authorities, after an explanation of why and how the knowledge would be used. The study was executed in accordance with the regulations regarding prior informed consent of the Code of Ethics of the Ethnobiological Society (International Society of Ethnobiology 2006). Two regions, Moyenne Guinée and Guinée Maritime were visited. All interviews were conducted in the mother tongues (Pular or Susu) of the participants in the study area.

The questionnaire consisted of 27 questions, for which participants provided the vernacular names of *Neocarya macrophylla* in their own languages, uses, part of the plant used, among other information (Appendix 2A). Following interviews, the researchers and local participants visited the field together to locate *Neocarya macrophylla*. One to six sites were visited per prefecture. Sites are defined as being a minimum of 3 km from other sites, or having a differentiating name from the community (see Table 1).

During the survey of trees we found that trees can be divided into two classes, those which have no, or very few (3-5) fruits, and those more mature, larger trees which have many dozens of fruits. These equate to those with a circumference of <11 cm (little or non-productive) or >11 cm (productive, fully mature) at 1.5 m above ground level, measuring with a circumference tape. Due to the large numbers of trees and limited time we did not count trees <11 cm, but estimate that overall, they were 3-4 times the number of the 780 trees counted >11 cm circumference. We also recorded 63 juveniles being below 1 m in height. For the 780 mature trees, we recorded detailed data for a subset 291 individuals (approx 2/5ths of the total) with up to 30 individuals sampled per subpopulation (depending on the number in the subpopulation) the circumference of up to four limbs per tree arising from ground level, calculating the average and from that the diameter at breast height. We also estimated the height of each tree to the nearest metre, its georeference with a Global Positioning System (Garmin), recording the habitat, substrate, type of threats, locality, date and making a herbarium voucher specimen of each tree of the subset (see data table Appendix 3). For each tree we collected and preserved a sample in silica gel for future genetic analysis, and c. 10 fruits per tree for subsequent chemical and nutritional analysis (to be the subject of future publication).

The collected voucher herbarium specimens were deposited at the National Herbarium of Guinea (HNG) and RBG Kew (K). The identification of the specimens was later confirmed by botanical experts from Kew and HNG.

### Distribution mapping

Georeferenced data for *Neocarya macrophylla* were downloaded from GBIF(gbif.org) supplemented by representative points for the study sites of this paper. All records complete with herbarium images were checked by the first author to avoid any previous misidentifications. Any specimens which had been misidentified were excluded from the distribution maps, and conservation assessment calculations. This resulted in cleaned and filtered matrix of 50 points (see Appendix 4) which was then used to generate distribution maps (Maps 1 & 2) and the conservation assessment (Appendix 5). Distribution maps were generated using QGIS v.3.14 (QGIS Development Team continuously updated).

### IUCN Red List conservation assessment

The global species conservation assessment for *Neocarya macrophylla* (Appendix 5) was produced following the guidelines set out in the IUCN Categories and Criteria v.3.1 (2012). To generate threat categories, the minimum Area of Occupancy (AOO) and estimated Extent of Occurrence (EOO) for each species was calculated using GeoCAT (Bachman *et al*. 2011). Data on threats in Guinea was based on the fieldwork for this study. For other countries we depended on the literature and from a colleague in Senegal.

### Species Conservation Action Plan

The methodology for developing the conservation action plan for *Neocarya* (Appendix 6) in this paper is based on the action plans developed by Couch *et al*. (2023) during the project “Towards a Red Data Book for Guinea.” The data included concerns the botanical description and ecology of the plant, distribution, phenology, use, conservation measures, threats, legislation. The information is also used to complement the IUCN Red List assessment.

## Results

### Field survey and interviews

#### Distribution in Guinea of the numbers of individuals of *Neocarya* recorded in the survey

The largest number of mature trees (above 11 cm circumference) of *Neocarya* recorded in the survey was in the Pita prefecture with 39% of the total recorded in the survey, closely followed by Telimélé and Gaoual prefectures. These three northern prefectures have more than 75% of the total mature individuals recorded in the survey of the seven prefectures surveyed for *Neocarya* (Figure 3).

**Fig. 1.**
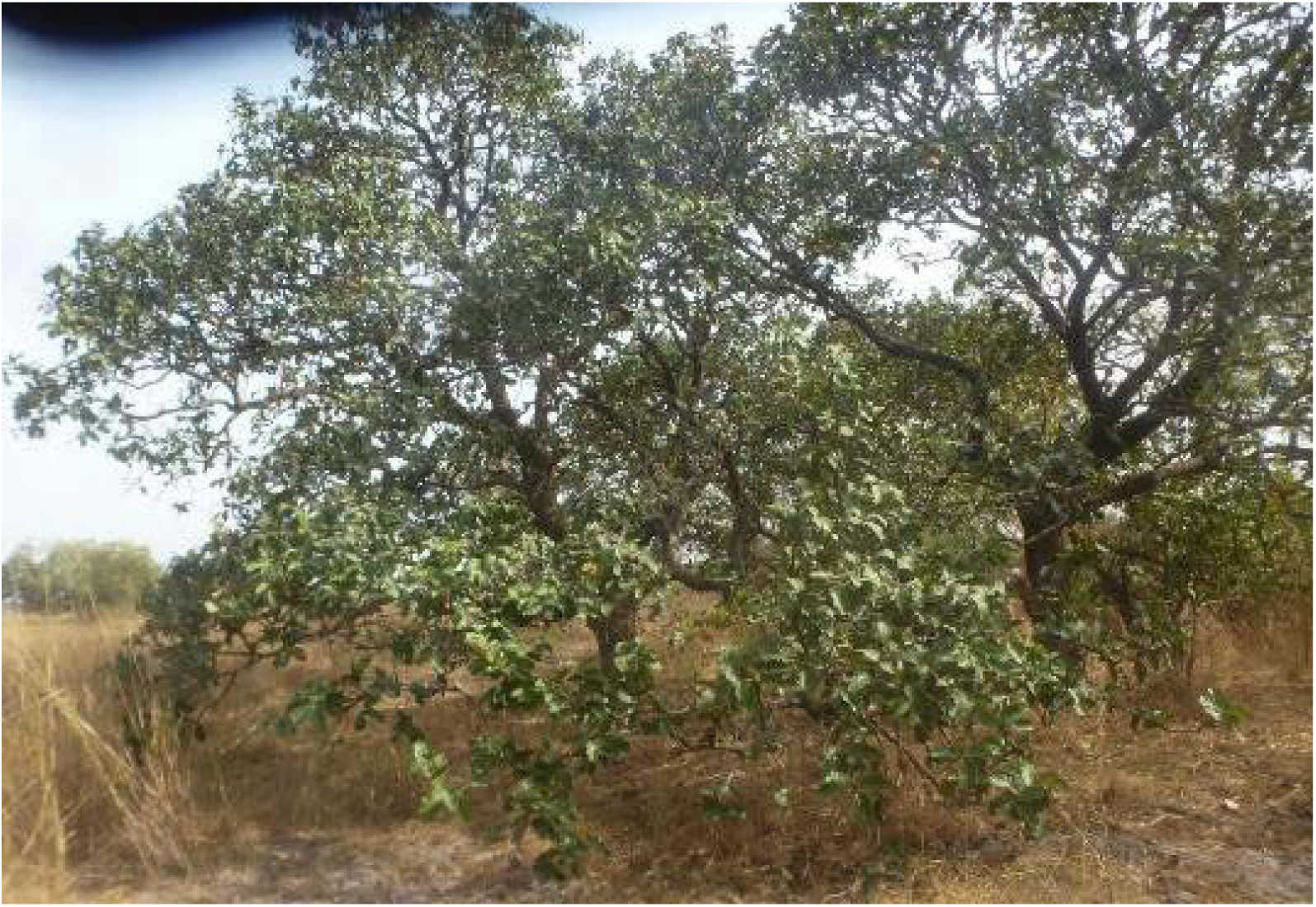
*Neocarya macrophylla* trees in habitat in Guinea. Photo Denise Molmou.

**Fig. 2.**
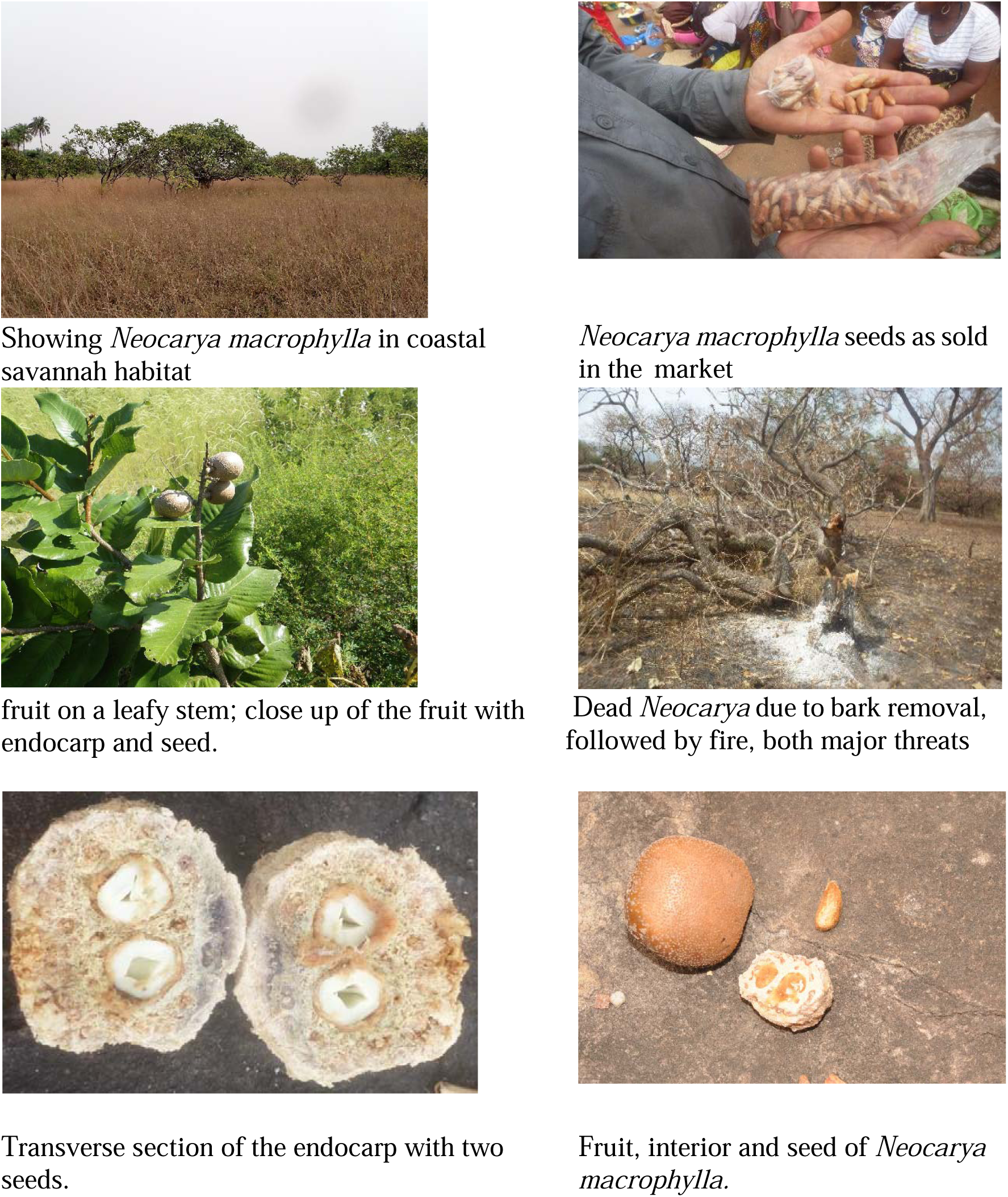
*Neocarya macrophylla* in habitat, markets, and threats. Photos Denise Molmou, Martin Cheek, Xander van der Burgt.

**Fig. 3.**
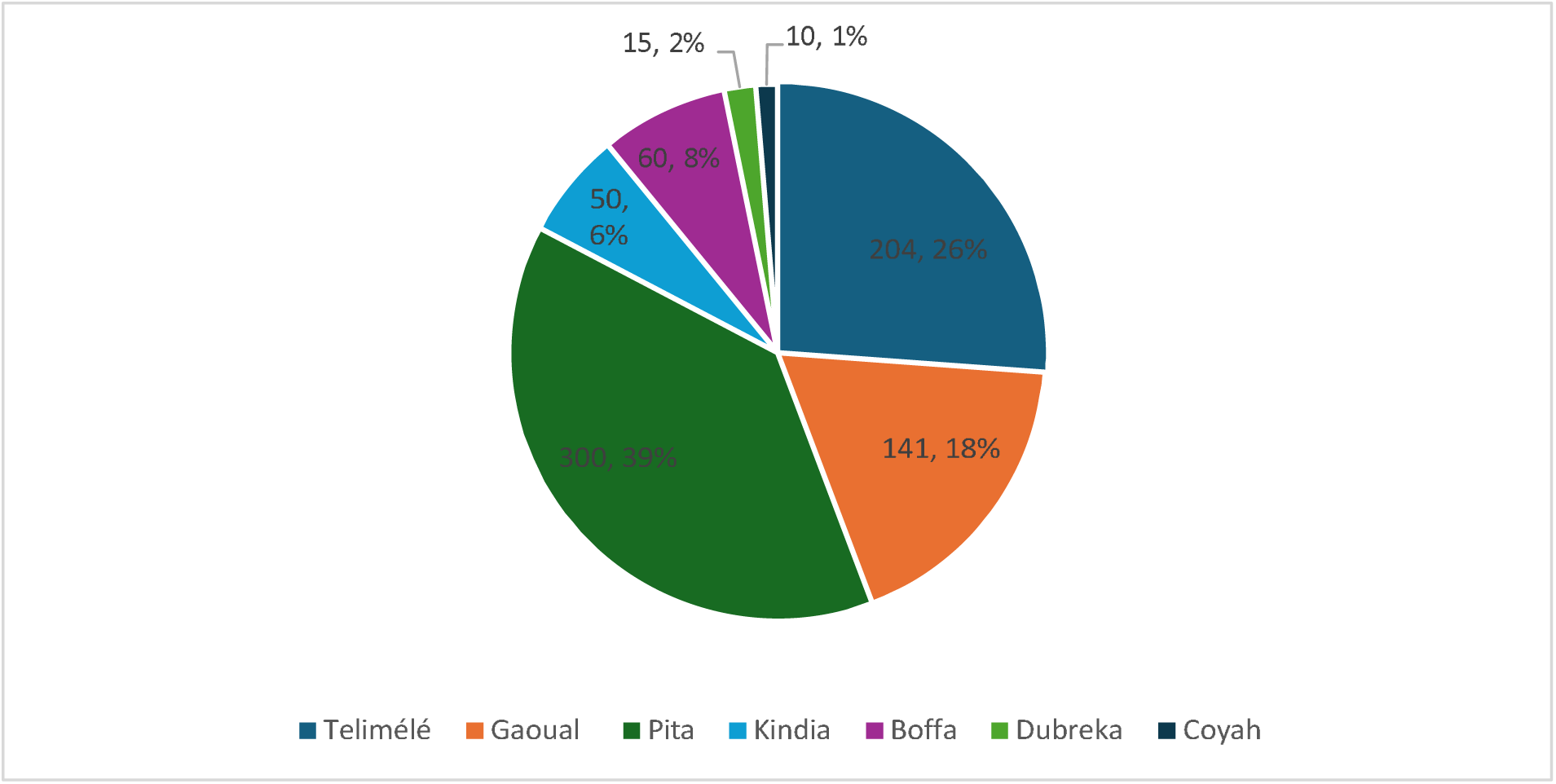
Number of mature individuals recorded of *Neocarya macrophylla* in the seven prefectures surveyed covering most of the prefectures thought to hold the species in Guinea.

##### Distribution of *Neocarya* by habitat type

Four main habitat types were identified for *Neocarya macrophylla* in the seven prefectures visited. The preferred habitat was shrub savannah on sandy soil (45% of individuals documented) and agricultural plain (33%), followed by bowal (18%) and wooded savannah (4%) (Figure 4). Trees in agricultural plains were not planted but have been allowed to persist since the original habitat, likely wooded savannah, was converted to agriculture. Shrub savannah is distinguished from wooded savannah by the lower height of the shrubs/trees (typically 2-5 metres) and lower density of woody plants (typically <10% cover vs 40 % cover). Bowal is typically treeless due to an impervious substrate of rock (e.g. sandstone) or concretised laterite (Couch *et al*. 2019b). Small trees can grow in this habitat where there are fissures allowing root penetration. All the juvenile plants inventoried were located at the edge of woodland and in clumps of *Neocarya* trees probably because there they have some protection from the pressure of human set bush fires each year.

**Fig. 4.**
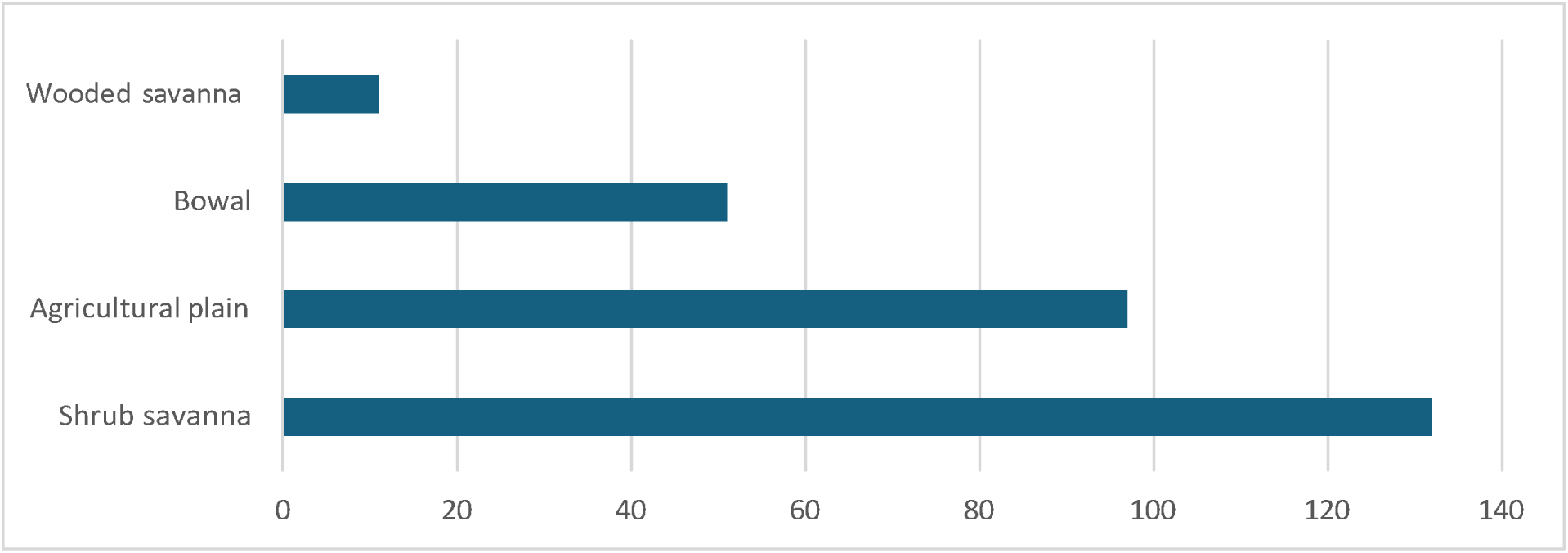
Number of *Neocarya macrophylla* individuals recorded per habitat type in the survey in Guinea.

##### Commerce and uses of *Neocarya* in the study area

Our survey in three prefectures (Télimélé, Gaoual, Pita) found that Bansouman or Koura-Bansouman (*Neocarya*) seeds are sold locally in plastic bags and in 250 g pots at a price of 5,000 GNF per container. This equates to 20,000 GNF or 2.3 USD per kg (current exchange rate 8,640 GNF = 1 USD in March 2025). For other areas (the prefectures of Kindia, Boffa, Dubreka, Coyah) price information was not available because the seeds are scarcer and more difficult to find in the markets in these prefectures. More extensive information on prices of *Neocarya* seeds at different points in the supply chain are given, mainly for Koundara prefecture, in the report of United Purpose (2023, Appendix 1). These show a great disparity between the price received per unit in the village of 4,600 GNF per unit at Dounia, and that paid in the market at Madina in Conakry of 35,000 GNF per unit. This equates to a mark up multiple of 7.6.

###### Uses

The fruit pulp (mesocarp) of the *Neocarya* fruit is consumed fresh in every prefecture of the study area and much appreciated, mainly used in the villages rather than sold in markets (Appendix 2). In one village, Ley Miro (Pita prefecture) we found that the mesocarp pulp is crushed in a machine and used to make a fruit juice that is consumed both locally and sold in the nearest market. Seeds or “nuts” of *Neocarya* (Bansouman or Koura-Bansouman) are also consumed as food either fresh or processed in every prefecture (Appendix 2). In the prefectures Kindia, Pita, Gaoual and Telimélé (Appendix 2) and also in Koundara (Appendix 1), but not in the remaining three prefectures of the survey (Boffa, Dubreka and Coyah) there are three other sets of usages:

1. The seeds are used to treat or to prevent arterial hypertension.
2. The powder from the bark of the plant is taken in lukewarm water to treat stomach aches.
3. The same mixture with cooking salt added is given to cattle to treat certain diseases, and also to aid fertility of the cattle (Appendix 2).

### Height, circumference, trunk number and substrate

For the 291 mature individuals (above 11 cm circumference) of the seven prefectures that were recorded in detail, the heights ranged from 3-15 m tall. Commonly (more than 50 % of individuals) trees had more than one trunk arising from ground level, but only nine individuals (c. 3%) were recorded with four trunks. The maximum circumference of a trunk at 1.5 m above ground level was 323 cm. All of the individuals occurred on sandy soils except for <10% where sandstone rocks were recorded as substrate (Appendix 3)

The most ubiquitous threat recorded to trees of *Neocarya* in the wild was human-set bush fires, commonly used in Guinea to provide fresh fodder for grazing animals, usually cattle, and to clear vegetation before cultivation. Trunks have thick bark and are fire resistant but are especially vulnerable if the bark has been removed for medicinal use, or if the fires are intense. Secondary threats were habitat clearance for sand quarrying and also clearance for agricultural land, both recorded as threats in two of the seven prefectures. Logging (the trees being cut for use of their wood either for carpentry or as firewood) was only recorded in one prefecture, Telimélé (Figure 5).

**Fig. 5.**
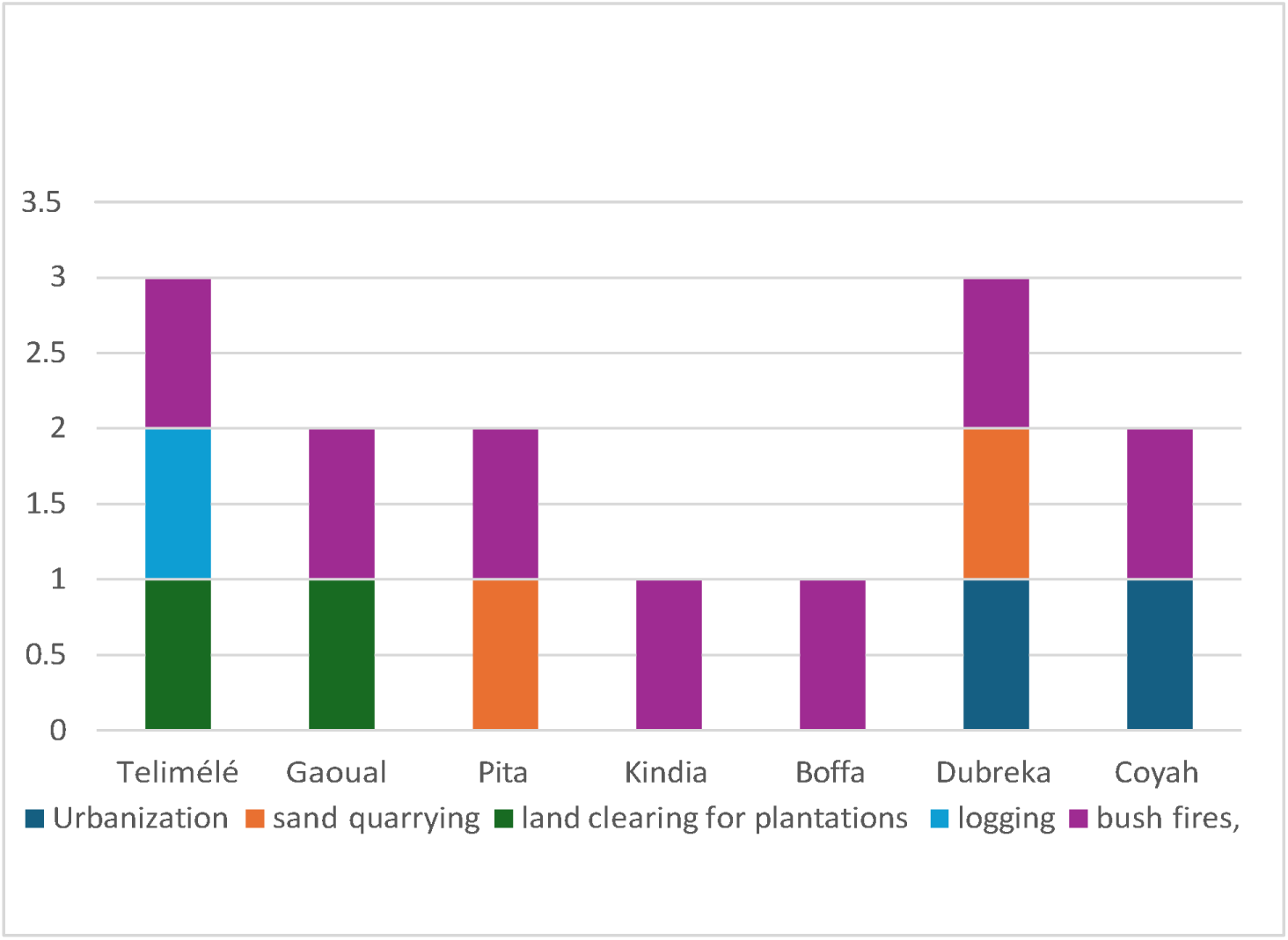
Analysis of the type, frequency and distribution of threats to *Neocarya macrophylla* observed in the field survey in Guinea.

### Distribution mapping

Based on the occurrence records from GBIF, georeferenced and vouchered herbarium specimens (HNG & K) and the field survey carried out for this study (the combined cleaned point data are at Appendix 4), the distribution of *Neocarya macrophylla* was mapped globally (Map 1), and for Guinea (Map 2).

### IUCN Red List conservation assessment

*Neocarya macrophylla* is a widespread tree species in West Africa, with some populations found in protected areas. It is a versatile tree that can be used for a variety of purposes, including wood, charcoal, food, fodder and medicines. Although the species is widespread and its estimated area of occurrence (EOO) is 2,927,233 km^2^, intensive and unsustainable harvesting (wood, charcoal) of this species has led to a reduction in its population and even total loss at some sites (at least at local or sub-population level in Guinea and Senegal), although no quantitative data are available. Threats are ongoing. Consequently, this species is assessed as Near Threatened (NT). We recommend monitoring sub-populations likely to be over-exploited or at risk to prevent an increase in the risk of extinction of this species in the future. See Appendix 5.

### Species Conservation Action Plan

The main threat for this species in the Maritime and Middle Guinea regions is uncontrolled bushfires, clearance for new agricultural land, sand quarrying, logging, and the use of its bark in traditional medicine. Pressure on *Neocarya macrophylla* habitat is high and in several places sites have been destroyed. No conservation action is in place for this species in Guinea, although village communities at several sites protect this species because its fruit is a source of nutrition and economic income for them. The species is reported to be cultivated in Panama (Prance & Sothers 2003) but otherwise is not known to be cultivated. It is known to be in the Millenium seed Bank but the 200 seeds, originating from Mali, have not been tested for viability (Duncan Sanders pers. comm. to Cheek, March 2025). It is possible that the seeds are recalcitrant due to their large size and their high lipid content. We observed in this study that seed germination from whole fruits works well, with good levels of germination and establishment of seedlings in July and August (mid wet season), while germination of extracted seeds seems to be less successful. See Appendix 6 for our Species Conservation Action Plan for *Neocarya macrophylla*.

## Discussion

Our study found that *Neocarya* trees are far more numerous in the prefectures at the western flank of the main Fouta-Djalon massif, that is the prefectures of Telimélé, Gaoual, Pita and Koundara, versus those of Kindia, Dubreka, Coyah and Boffa (Fig. 3). We deduce that numbers are lower in the Kindia area partly because of the prevailing granite outcrops in that area which appear unsuitable for *Neocarya*, and are lower in Dubreka, Coyah and Boffa areas because so much of the habitat has been converted into areas of habitation or has been degraded by human impacts as these prefectures are much more densely populated with humans than the first mentioned. There may be other explanatory factors such as differences in climate but we have not been able to investigate that in this study.

In terms of population structure, our results for Guinea were loosely similar to those for Niger (Guimbo *et al*. 2016a) in that numbers in the smallest size class (taken in our study as <11 cm circumf., and in Niger as the 0-4 m height class so not directly comparable) comprise 2/3 to ¾ of the total number of individuals.

The differences in numbers of individuals between different habitats is striking (Fig. 4). The largest number of trees was found in shrub savanna on sandy soils which appears to be the habitat most favoured by the species in Guinea. This habitat, having shallow sandy soils above rock, is far less impacted by agricultural clearance being less amenable to cultivation than wooded savannah which has deeper, more cultivable soils. Due to these deeper soils, wooded savanna, in the neighbourhood of villages has been largely converted to ‘agricultural plain’, which approaches shrub savanna in terms of the total numbers of mature individuals recorded. The low numbers we recorded in woodland savanna may not be because that habitat is unsuitable, but because it is so scarce near villages, having been converted to fields. The low numbers of trees in bowal habitat is to be expected because that habitat with solid surface substrate, has few and sparse woody plants, by definition. We have found no studies in other countries for this species in which partition by habitat is documented as we have. The only habitat referred to in such studies is savanna. This may be because habitat diversity is lower elsewhere within the range of the species, e.g. bowal habitat is much less frequent elsewhere in West Africa. Since we did not record population densities per se, we cannot compare densities with those studies that record this e.g. in Niger.

In terms of the uses of the species, within Guinea our data shows a great difference between the prefectures of Kindia, Boffa, Coyah and Dubreka where we did not record as we did in the other prefectures, the medicinal uses of the seed for hypertension, nor the bark for stomach ache, and for treating illnesses in cattle. This may be explained by the fact that for Boffa, Coyah and Dubreka prefectures we had difficulty in finding people to interview on the uses of this species, and those few we did find did not have the traditional knowledge needed. Possibly, with more time we might have located such people and found that the same uses obtain elsewhere. However, in Kindia there seems to be a genuine difference in the traditional use of this species from all the other prefectures studied. This is because although the medicinal uses referred to above were unknown among the many people we interviewed, a new and unique use for the species was recorded (treating intestinal parasites of children using the endocarp hairs embedded in banana as in *Bafodeya benna* (Scott Elliot) Prance ex F.White (Cheek pers. obs. 2011). Also in Kindia, the local name (Gnamouie) is not used elsewhere in our study (see Table 1).

In other countries within the range of the species, the fruits and the seeds are used as drink and food including seed oil in the same way as reported in our study e.g. in Guimbo *et al*. (2017) and in Yusuf *et al*. (2024). However, elsewhere, in Nigeria, additionally, the fruits are used to make a tangy tasting gruel by pounding in water and thickening with maize or cassava flour, and the fruit juice is also concentrated to make a fragrant syrup (Yusuf *et al*. 2024).

In terms medicinal uses in other countries, the uses in Guinea that we report in this paper are also used elsewhere e.g. Nigeria and Niger (Datti *et al*. 2020, Guimbo *et al*. 2017, Yusuf *et al*. 2024). However, again additional uses are reported e.g. in Niger the bark and root decoction is additionally used to treat haemorrhoids, and the leaves, mixed with those of other species are used as a treatment against diarrhoea in small children (Guimbo *et al*. 2017). In Northern Nigeria additional medicinal uses to those we recorded in Guinea are in treatment of asthma, skin infections, pulmonary troubles, ear and eye infections (e.g. conjunctivitis), snake bites and cancer (Yusuf *et al*. 2024).

In Nigeria and in Niger, several uses unknown in Guinea apply, including as soap (likely manufactured from the seed oil), as dye, glue, as a termite repellant, and for fabricating household utensils such as pestles and mortars (Guimbo *et al*. 2016b, Yusuf *et al*. 2024). The geographic variation in threats to *Neocarya* within Guinea as reported in our study partly reflects the differences in human population density which being higher in Boffa and Dubreka prefectures in study sites for the species, necessarily results in urbanization being recorded as a threat there and not in other, less densely populated prefectures. On the other hand, although Gaoual and Telimélé prefectures are more sparsely populated they are more agricultural, explaining the higher incidence of agricultural clearance as a threat in those study sites.

Threats to *Neocarya* in other countries differ in level, and even in nature from those we recorded in Guinea. While fire is the universal threat in Guinea at all study sites, this appears not to be a major threat in other countries such as Niger, where overharvesting of the wood for making tools (e.g. pestles and mortars) and as fuelwood is the major threat (Kolofane *et al*. 2018b, Guimbo *et al*. 2016b). In contrast logging of trees was only a threat in one of the seven prefectures in our study. The secondary threat in Niger is overharvesting of the bark for making rope and medicine, which by itself is not one of the top five threats in Guinea (which after fire are urbanization, sand quarrying, clearance for agriculture and lowest, logging see Fig. 5). Desiccation and impoverishment of soils, a threat unreported in Guinea, is stated as a major threat in Niger (Kolofane *et al*. 2018b).

## Conclusions

Our field study of *Neocarya macrophylla* in the wild in Guinea covered villager interviews and tree surveys in seven of the ten prefectures where it is known nationally. We conclude that the species is understudied, underutilised and under-appreciated in Guinea. It is important for local populations above all in Moyenne Guinea as an Indigenous Fruit Tree which produces both delicious edible fruit and valuable seeds (’nuts’) of large size and good flavour, important nutritionally especially in the months of January to March. The seeds are a source of protein (c. 20%,) and as a source of oil, and the species is also is a source of medicinal treatment for hypertension (the seeds) and for treating dysentery and diarrhoea in humans, and also illnesses and infertility in cattle (the bark). The leaves also have medicinal uses, as do the endocarp hairs for treating internal parasites in children. Evidence that the species is understudied is that ours is the first study focussed on *Neocarya macrophylla* in Guinea. In contrast, in Niger, 12 papers on this species as a useful species can be found (see Introduction). *Neocarya macrophylla* has not been included in publications on the priority indigenous fruit trees of Guinea, (Camara *et al*. 2002, N’Diaye *et al*. 2003). This may partly be explained by the remoteness of the four most productive prefectures for *Neocarya* from the capital, Conakry and from other major cities. Again, over the border in Sierra Leone where the species is also not among the priority Indigenous Fruit Trees documented (Jusu & Cuni-Sanchez 2017). Evidence of under-utilisation is that in Niger and Nigeria, the species is used to manufacture numerous products, which uses were not recorded in Guinea our interviews in this study (see Discussion above).

The large differential between the price paid per unit of seeds in Conakry and that in the village, a multiple of 7.3, and that raw, unprocessed seeds are exported to other countries as far as Nigeria suggests also suggests that the seeds are under-valued and under-exploited in Guinea. That fruits are often left unharvested confirms this, and suggests that the value chain does not deliver sufficient return to wild harvesters considering the high difficulty of seed extraction. If these issues were addressed, trees in the wild might be better protected from the numerous threats we document which have resulted in a global conservation assessment of Near Threatened and necessitated a Conservation Action Plan (both this paper).

Future work

1. As significant differences were reported in seed nutrition and chemistry between Niger and the productive prefectures in Guinea (a distance of c. 1700 km as measured on Google Earth Pro), the data characterising the species in Niger may not be fully applicable to Guinea so detailed characterisation of the Guinea subpopulations is advisable. Recording densities of trees, and calculation of fruit yield per tree in Guinea are advisable.
2. Measurement of age taken from germination to fruiting and also the productive longevity of trees in production (for how many years do they continue to bear useful crops) would give useful information in managing populations in Guinea.
3. Mapping areas of trees to quantify potential production, perhaps using near infra red sensors from drones to identify the species more efficiently.
4. Field trials of planted trees would enable selection of elite individuals more multiplication, e.g. those with heavier fruit yields, productive in early years.
5. Development of a machine to extract seeds efficiently, quickly, and more safely from the endocarps than with a machete. This would remove a major constraint in production.
6. Development in Guinea of novel high value products following the Nigerian example (e.g. manufacture of soaps and dyes).

## Acknowledgements

This paper was founded on two previous projects in Guinea, one funded by the European Union through a Global Biodiversity Information Facility - Biodiversity Information for Development (GBIF-BID) grant and the other by the UK government through a Darwin Initiative grant in 2016–2019). The Ellis Goodman Family Foundation are thanked for the initial funding to the first author that enabled her to be registered as a Doctoral student at the University of Ghent, Belgium. This paper constitutes a chapter for her doctoral thesis. The Herbier National de Guinee, IREG and Universite Gamal Abdel Nasser, Conakry is thanked for providing the field permits, logistic support and for supporting the first author for many years of her doctoral studies. Rio Tinto Guinea Biodiversity Dept. sponsored study visits of the first author to RBG, Kew in 2024 and 2025 when this paper was written up. The first author especially thanks Harry Nevard of Rio Tinto for support.

We thank Professor Ben Bennett of Natural Resources Institute University of Greenwich for highly useful comments on our study. Two anonymous reviewers are thanked for comments on an earlier version of this manuscript.

## Confict of Interest

The authors declare that they have no conflict of interest.

## Notes

### Competing Interest Statement

The authors have declared no competing interest.

