## Supplemental file 2A for "Traditional uses, population, threats and conservation of the bansouman or gingerbread plum *Neocarya macrophylla* (Chrysobalanaceae) in Republic of Guinea (West Africa)"

**Appendix 1.**

Example of a completed interview form used for the villagers on *Neocarya macrophylla* (the responses are a composite from more than one village in Telimélé & Gaoual prefectures, translated into English;).

1. Do you know *Neocarya* *macrophylla*? Yes

2. What is its local name? Bansouman

3. Does it grow in the vicinity of the village?

- Not only in the vicinity of the village, it grows everywhere according to the needs of its habitat

4. What habitat does it grow in? Forest, alongside fields?

- The plant grows in wooded savannahs, on the plains, at the edge of rivers on sandy soils sometimes rocky.

5. Is it cut down when land is cleared for crops? Or is it selectively kept?

- Agriculture is not generally possible on the habitat of this plant, thirsty on the plains where the plant grows with space, the sofa is open, the dead leaves enrich the soil. No foot is cut for agriculture. Confirmed by a citizen of kithiar

6. Is it ever planted? Do you know whether anyone has tried to grow it?

- It’s a wild plant, we have not been shown anyone who has planted in a place.

7. Do you use it? Yes

8. What do you use it for?

- It is used for food and for medicine.

9. What parts of the plants do you use? Nuts, Fruit peel

10. Do the parts of the plant have different names? no, the - - common name is bansouman

11. How much do you appreciate the nuts?

- - Nuts very shower, we bite like peanuts, and they fight against blood pressure.

12. Are the nuts a key part of your diet at a particular time of the year? ON

13. Do you collect the nuts?

Yes

14. What months do you harvest the nuts? - January February March,

15. Do you know anyone in the village who collects the nut?

16. Who collects the nuts? Individuals (male or female) or group activity?

- In general, women and children. Have to harvest for remembrance of our small needs if not, the job of harvesting nuts is very difficult, and the economic return is very low. Confirms a citizen in Faro.

17. How many nuts are collected by you? The village?

- No idea in the entire survey area

18. How do you take the nuts out of the fruit?

- - We cut the fruits to obtain the nuts with a cutter or machete.
- - The nuts are stored in a bag or in a tightly closed jar and keep in a dry place.

19. How much time does this take?

- nuts can be kept all year round.

20. Do you sell it?

- Yes

21. How much money do you get for 1kg of nuts?

- If the small pot is 5000 fg, then the kilo is 20000fg.

22. Is it sold locally? Or does it go somewhere else?

- - We sell it locally, but sometimes people come by order and sent.

23. Do you store the nuts? How? Where?

- - Yes, we can store the nuts in a fabric bag to avoid the heat or in a jar tightly closed and keep in a dry place.

24. Would you be interested in harvesting the trees if the process to extract the nuts is made shorter?

- We will be more motivated if the extradition method becomes easier.

25. Would you be interested in harvesting the trees if the price you receive is higher?

Yes

26. If the price is better or there is more demand, will that stop people from clearing habitat where the trees occur?

- Yes, if it becomes an important economic source, we will take care to keep it jealously

27. How would this affect your livelihood?

- we think it will affect positively,
