## Supplemental file 2B for "Traditional uses, population, threats and conservation of the bansouman or gingerbread plum *Neocarya macrophylla* (Chrysobalanaceae) in Republic of Guinea (West Africa)"

Appendix X Uses of *Neocarya macrophylla*. Summaries by prefecture of the results of the 18 village questionnaires from our study:

| Prefecture | Uses |
| --- | --- |
| Kindia | The seeds of Gnamouie or Bansouma are used by the interviewees as a medicine for treating or preventing arterial hypertension. The powdered bark of the plant, mixed with cooking salt, is given to cattle not only to treat certain illnesses, but also to promote fertility. The decoction of leaves and roots is used against dysentery and diarrhea. The fruit is edible, and the hairs from the inner surface of the endocarp are eaten with bananas to combat parasites in children. |
| Pita | The seeds of Bansouma are eaten to treat or prevent arterial hypertension, and diabetes, the powder of the bark of the plant is taken in tepid water to treat stomach ache (the dysentery and the diarrhoea) the same mixture with kitchen salt is given to oxen not only to treat certain diseases, but also facilitates fecundity.  The fruit flesh is edible with a pleasant taste and can be made into fruit juice for consumption. |
| Gaoual | The seeds of Koura- Bansouma are eaten raw to treat hypertension, diabetes, dysentery and diarrhoea. A decoction of the leaves or roots is taken as a mouthwash against tooth decay.  The fleshy fruit is edible, and the seeds contain oil that can be used in cooking.  Fallen leaves make the soil fertile for farming. |
| Télimélé | The seeds of Bansouma are eaten to treat or prevent arterial hypertension, the powder of the bark of the plant is taken in tepid water to treat stomach aches, the same mixture with cooking salt is given to oxen not only to treat certain diseases, but also facilitates fertility, the fruit flesh is edible, the seeds are eaten raw, and can be roasted to eat, dishes of the roasted seed are prepared as food. |
| Boffa | The flesh fruit is edible with a sweet taste, and the seeds are eaten as a medicine to combat high blood pressure. |
| Dubréka | The fruit flesh is edible also the seeds. |
| Coyah | Trees are protected because they give shade. Fruits and seeds are edible. |
