## Supplemental file 5 for "Traditional uses, population, threats and conservation of the bansouman or gingerbread plum *Neocarya macrophylla* (Chrysobalanaceae) in Republic of Guinea (West Africa)"

**Draft**

**Neocarya macrophylla - (Sabine) Prance ex F.White**

PLANTAE - TRACHEOPHYTA - MAGNOLIOPSIDA - MALPIGHIALES - CHRYSOBALANACEAE - Neocarya - macrophylla

**Common Names:** Banssouman (Susu), Koura Bansouman (Susu)
**Synonyms:** Parinari macrophylla Sabine

**Red List Status**

NT, (IUCN version 3.1)

### Red List Assessment

#### Assessment Information

Date of Assessment: 2025-02-18

| **Reviewed?** | **Date of Review:** | **Status:** | **Reasons for Rejection:** | **Improvements Needed:** |
| --- | --- | --- | --- | --- |
| true | 2025-02-24 | Passed | - | - |

Assessor(s): Molmou, D.

Reviewer(s): Diop, F.N.

Contributor(s): Cheek, M.

Facilitators/Compilers: Couch, C. & Bevan, H.

Institution(s): IUCN SSC West Africa Plant Red List Authority & Royal Botanic Gardens, Kew

Regions: Global

#### Assessment Rationale

Neocarya macrophylla is a tree species that is widespread in West Africa, and some populations are found in protected areas. It is a versatile tree that can be used for a variety of purposes, including wood, charcoal, food, fodder, and medicine.  Although the species is widespread and its estimated area of occurrence (EOO) is 2,927232,956 km2, intensive and unsustainable harvesting (wood, charcoal) of this species has resulted in a reduction in its population (at least at the local or subpopulation level), although no quantitative data are available. The threats are ongoing. Therefore, this species is assessed as Near Threatened (NT). We recommend monitoring of potentially exploitable subpopulations to prevent the increased risk of extinction of this species in the future.

### Distribution

#### Geographic Range

This species is native to West Africa. It has a wide geographical distribution in West Africa: Senegal to Nigeria.

The size of the range and the number of locations would increase significantly with more survey work, as a common species. Neocarya macrophylla is under-represented by herbarium specimens.

The author observed in Guinea that the species has been lost at some sites because of the threats mentioned.

Specimens recorded in the Pic de Fon classified forest in Guinea and in the Moyen Bafing National Park have not been confirmed. The identification was probably mistaken for another species.

Some sources (GBIF 2025, POWO 2025) indicate that the species is found in South Sudan, Cameroon and Côte d'Ivoire, but we have not found any evidence to support this claim.

#### Area of Occupancy (AOO)

| **Estimated area of occupancy (AOO) - in km2** | **Justification** |
| --- | --- |
| 176.000 | Calculated using GeoCAT (Bachman et al. 2011). |

| **Continuing decline in area of occupancy (AOO)** | **Qualifier** | **Justification** |
| --- | --- | --- |
| Yes | Inferred | The author observed in Guinea that the species has been lost at some sites due to the threats mentioned and this is supposed to continue. |

#### Extent of Occurrence (EOO)

| **Estimated extent of occurrence (EOO)- in km2** | **EOO estimate calculated from Minimum Convex Polygon** | **Justification** |
| --- | --- | --- |
| 2927233 | true | Calculated using GeoCAT (Bachman et al. 2011). |

| **Continuing decline in extent of occurrence (EOO)** | **Qualifier** | **Justification** |
| --- | --- | --- |
| Unknown | - | - |

#### Locations Information

| **Number of Locations** | **Justification** |
| --- | --- |
| 44 | Based on the cleaned file of data points used for GeoCAT in this assessment (see attached). |

| **Continuing decline in number of locations** | **Qualifier** | **Justification** |
| --- | --- | --- |
| Unknown | Inferred | - |

| **Extreme fluctuations in the number of locations** | **Justification** |
| --- | --- |
| No | - |

#### Very restricted AOO or number of locations (triggers VU D2)

| **Very restricted in area of occupancy (AOO) and/or # of locations** | **Justification** |
| --- | --- |
| No | - |

#### Elevation / Depth / Depth Zones

Elevation Lower Limit (in metres above sea level): 10

Elevation Upper Limit (in metres above sea level): 1200

#### Map Status

| **Map Status** | **Use map from previous assessment** | **How the map was created, including data sources/methods used:** | **Please state reason for map not available:** | **Data Sensitive?** | **Justification** | **Geographic range this applies to:** | **Date restriction imposed:** |
| --- | --- | --- | --- | --- | --- | --- | --- |
| Done | - | We used cleaned GBIF data and field samples. | - | - | - | - | - |

#### Biogeographic Realms

Biogeographic Realm: Afrotropical

### Occurrence

#### Countries of Occurrence

| **Country** | **Presence** | **Origin** | **Formerly Bred** | **Seasonality** |
| --- | --- | --- | --- | --- |
| Benin | Extant | Native | - | Resident |
| Gambia | Extant | Native | - | Resident |
| Guinea | Extant | Native | - | Resident |
| Guinea-Bissau | Extant | Native | - | Resident |
| Liberia | Extant | Native | - | Resident |
| Mali | Extant | Native | - | Resident |
| Niger | Extant | Native | - | Resident |
| Nigeria | Extant | Native | - | Resident |
| Senegal | Extant | Native | - | Resident |
| Sierra Leone | Extant | Native | - | Resident |
| Togo | Extant | Native | - | Resident |

### Population

- There is no information on the population size or current trends of this species globally.

The Guinea subpopulation is in considerable decline, mainly due to unsustainable overexploitation for service and medicinal timber combined with reported intensive fires in its habitat (Per. obs. Molmou 2020-2024).

#### Population Information

Current Population Trend: Unknown

### Habitats and Ecology

Neocarya macrophylla occurs in a range of forest habitats but remains a characteristic savannah species. It is found in forest edges, open forests, open ground, coastal lowlands. Up to 12 m tall, this species usually flowers during the rainy season and the fruits are often present at the beginning of the dry season. The spread of seeds is probably due to animals and sometimes even to people.

#### IUCN Habitats Classification Scheme

| **Habitat** | **Season** | **Suitability** | **Major Importance?** |
| --- | --- | --- | --- |
| 2.1. Savanna -> Savanna - Dry | - | Suitable | Yes |
| 2.2. Savanna -> Savanna - Moist | Resident | Suitable | Yes |

#### Continuing Decline in Habitat

| **Continuing decline in area, extent and/or quality of habitat?** | **Qualifier** | **Justification** |
| --- | --- | --- |
| Yes | Inferred | Frequent fires started by herders degrade the habitat of this species throughout its range and in particular in Guinea. |

#### Systems

System: Terrestrial

#### Plant / Fungi Specific

Wild relative of a crop? No

**Plant and Fungal Growth Forms**

Tree - large

### Use and Trade

#### General Use and Trade Information

This species has multiple uses, including food, medicine, and energy (firewood). The fruit pulp and seeds are edible. They are highly valued by rural communities who harvest it in the wild It is often sold in local markets and is kept and available all year round. The plant also has a range of medicinal applications. The seeds are eaten raw, or ground and used as a sauce for condiments, or used for traditional oil extraction.

The roots, leaves, and bark are also used in traditional pharmacopoeia. A decoction of roots, leaves and bark is used to treat sprains and fractures and to combat stomach aches, dysentery, blood pressure and haemorrhoids.

| **Subsistence:** | **Rationale:** | **Local Commercial:** | **Further detail including information on economic value if available:** |
| --- | --- | --- | --- |
| Yes | Fruits and nuts are edible, organs such as roots, leaves, and bark also used in traditional pharmacopoeia | Yes | - |

National Commercial Value: Yes

| **End Use** | **Subsistence** | **National** | **International** | **Other (please specify)** |
| --- | --- | --- | --- | --- |
| 1. Food - human | true | true | true | - |
| 2. Food - animal | true | true | - | - |
| 3. Medicine - human & veterinary | true | true | - | - |
| 7. Fuels | true | true | - | - |
| 11. Other household goods | true | true | - | - |

Is there harvest from captive/cultivated sources of this species? No

Trend in level of total offtake from wild sources: Increasing

Harvest Trend Comments: Removing bark from the stem, cutting roots for medicinal uses, picking up all fruit from under the foot for consumption

### Threats

The greatest threat to this species in Guinea is uncontrolled bush fires in its wooded habitat (savannah), logging and also the use of its bark in traditional medicine. Fire pressure on Neocarya macrophylla habitats is high. The Bowé, montane and submontane grasslands are the most affected by bushfires in Middle Guinea and Maritime Guinea. Pastoralists, by frequently setting up bush fires in the savannahs to obtain fresh growth of plants serving as fodder for their animals, destroy the habitats of the species.

A study in Niger also indicates that overexploitation and adverse weather conditions are potential threats to this species. In addition, erosion contributes to the uprooting of plant (Aboubacar et al. 2018)

In both countries, the use of firewood significantly reduces populations.

In Senegal, the exploitation of fruits, roots and bark for food and medicinal uses is a major threat to local populations (Fatimata Diop Niang pers. obs. to Molmou 2025).

#### Threats Classification Scheme

| **Threat** | **Timing** | **Scope** | **Severity** |
| --- | --- | --- | --- |
| 2.1.1. Agriculture & aquaculture -> Annual & perennial non-timber crops -> Shifting agriculture | Ongoing | Majority (50-90%) | Slow, Significant Declines |
| 2.3.1. Agriculture & aquaculture -> Livestock farming & ranching -> Nomadic grazing | Ongoing | Majority (50-90%) | Rapid Declines |
| 5.2.1. Biological resource use -> Gathering terrestrial plants -> Intentional use (species is the target) | Ongoing | Majority (50-90%) | Slow, Significant Declines |
| 5.3.3. Biological resource use -> Logging & wood harvesting -> Unintentional effects: (subsistence/small scale) [harvest] | Ongoing | Majority (50-90%) | Slow, Significant Declines |
| 7.1.1. Natural system modifications -> Fire & fire suppression -> Increase in fire frequency/intensity | Ongoing | Majority (50-90%) | Rapid Declines |

### Conservation

Neocarya macrophylla has been reported in many protected areas (including classified forests) such as Senegal (Niokolo-Koba National Park, Lower Casamance National Park), Guinea, in the classified forests of (Fello Sounga, Sala, Gangan), Nigeria in the forest reserve of (Cross River North), Sierra Leone (Western Peninsula National Park). It is recommended that subpopulations of this species be monitored and that sustainable harvesting measures be adopted for bark.

A study conducted in Benin found good regeneration of this species by seed germination (Aboubacar et al. 2014). Seeds have also been sown in wild areas (Aboubacar et al. 2018). It is also recommended to set up firebreak networks to protect survivors (D. Molmou pers. obs. 2025).

#### Conservation Actions In- Place

| **Action Recovery Plan** | **Note** |
| --- | --- |
| Yes | An action plan for the conservation of Neocarya macrophylla will soon be published |

| **Systematic monitoring scheme** | **Note** |
| --- | --- |
| Yes | - |

| **Conservation sites identified** | **Note** |
| --- | --- |
| Yes, over part of range | Mount Gangan and Kounounkan are potential national parks proposed to the national park network (World Bank and Government of Guinea, 2023). |

| **Area based regional management plan** | **Note** |
| --- | --- |
| Yes | - |

| **Invasive species control or prevention** | **Note** |
| --- | --- |
| Not Applicable |  |

| **Harvest management plan** | **Note** |
| --- | --- |
| No | No known harvest management plan. |

| **Successfully reintroduced or introduced benignly** | **Note** |
| --- | --- |
| No | - |

| **Subject to ex-situ conservation** | **Note** |
| --- | --- |
| Yes | Propagation and sharing of seeds or plants in botanical gardens |

#### Important Conservation Actions Needed

| **Conservation Actions** | **Note** |
| --- | --- |
| 3.1.3. Species management -> Species management -> Limiting population growth | - |
| 6.2. Livelihood, economic & other incentives -> Substitution | - |

#### Research Needed

| **Research** | **Note** |
| --- | --- |
| 2.2. Conservation Planning -> Area-based Management Plan | - |
| 3.2. Monitoring -> Harvest level trends | - |
| 3.4. Monitoring -> Habitat trends | - |

### Ecosystem Services

#### Ecosystem Services Provided by the Species

|  | **Importance:** | **Geographic range of benefit:** |
| --- | --- | --- |
| 3. Flood Control | 3 - Some Importance | Local |
| 8. Habitat Maintenance | 2 - Important | Local |
