## Supplemental file 6 for "Traditional uses, population, threats and conservation of the bansouman or gingerbread plum *Neocarya macrophylla* (Chrysobalanaceae) in Republic of Guinea (West Africa)"

Conservation d’Action Plan (PAC)

*Neocarya macrophylla* (Sabine) Prance (Chrysobalanaceae)

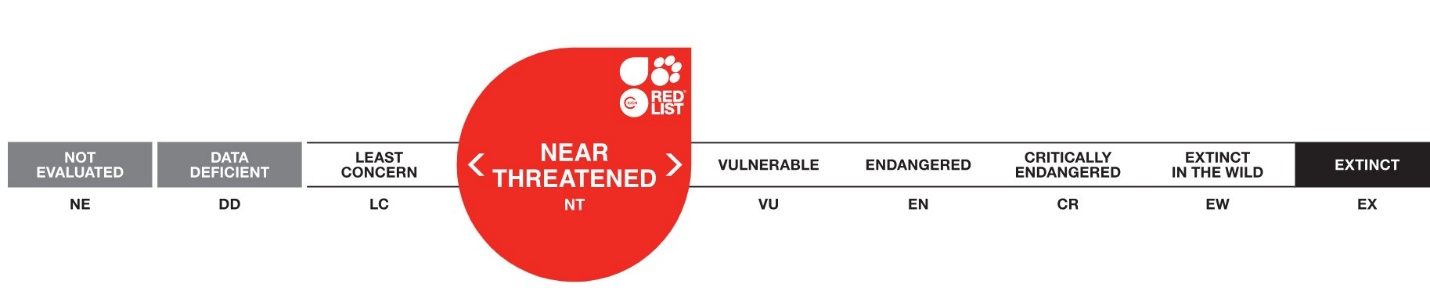

### Status, Description, habitat, and ecology:

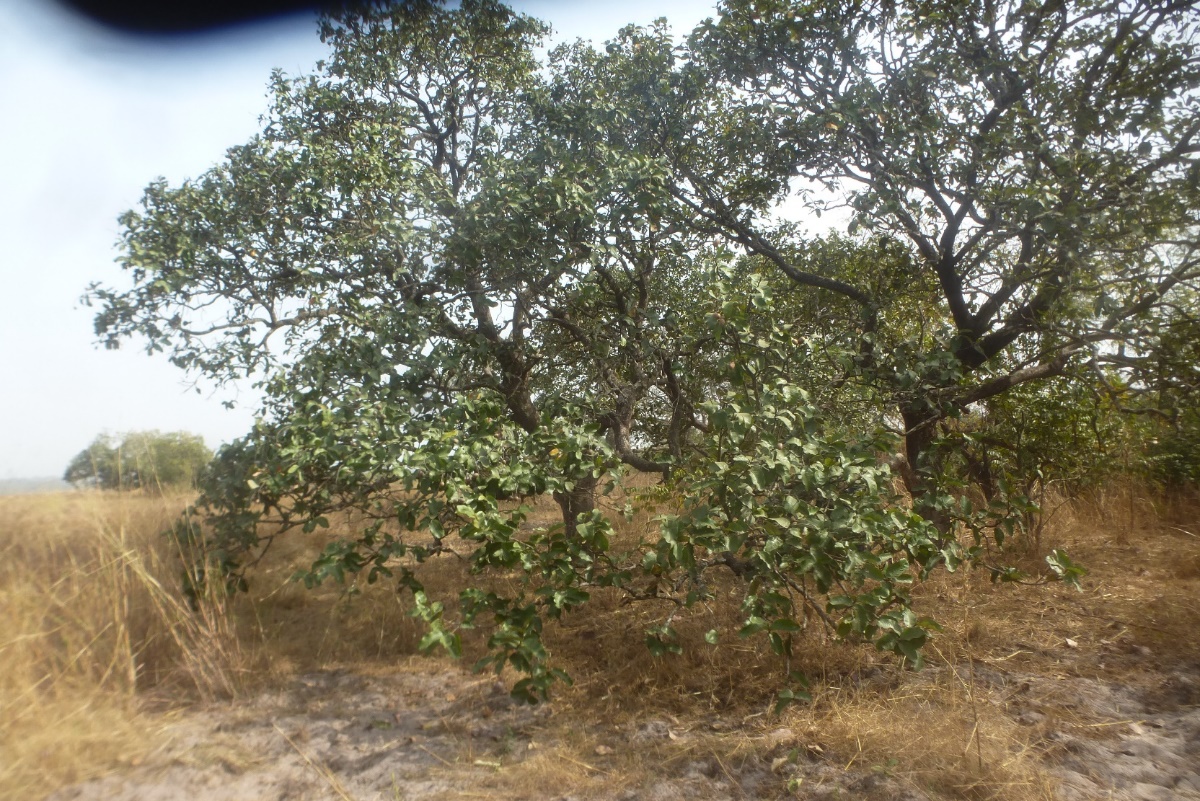
Savanna tree, reaching 12 m in height; young branches densely brown tomentose. Leaves alternate, simple, short-stalked; leaf blade ovate or elliptical, 10 to 25 cm long and 5 to 15 cm wide, rounded to subcordate at the base, rounded to slightly acuminate at the apex, densely white tomentose beneath, pinnate, with 15 to 20 pairs of lateral veins. Flowers small, white, or pinkish white, grouped in narrow, little-branched panicles; receptacle shorter than sepals; stamens fertile, 15. Fruits fleshy, drupaceous, ellipsoid, approx. 5 cm long, rough, edible (Lisowski 2009).

**General distribution**: West tropical Africa. The plant is widely distributed along coastal savannas from Senegal, Gambia, Guinea-Bissau, Guinea, Sierra Leone to Liberia, and also inland wooded savannas of southern Mali, Niger and northern Nigeria (Prance & Sothers 2003).

**Distribution in Guinea**: Boffa region, around Mankountan, Songolon riverbank, near Boffa Boké road, *Lisowski* 51472; Dubréka, *Lisowski* 51306; Kindia region, N of Foulaya, *Lisowski* B 7061; ibid, below Barrage Kalé, (Lisowski 2009). It is most frequent in Koundara, Pita and Télimélé Regions (Molmou et al. this paper).

**Habitat and ecology** Neocarya macrophylla occurs in a range of habitats but is a characteristic savannah species of sandy soils. It is found in shrub savannah, tree savannah, open woodland, forest edges, open ground, coastal lowland savannah. Rare on sandstone hillsides and granite rocks. (Prance & Sothers 2003; Yusuf et al. 2024). Seed distribution is probably by animals and even people.

**Phenology: Flowering**: May - December. **Fruits**: January – March

**Population status – current research**
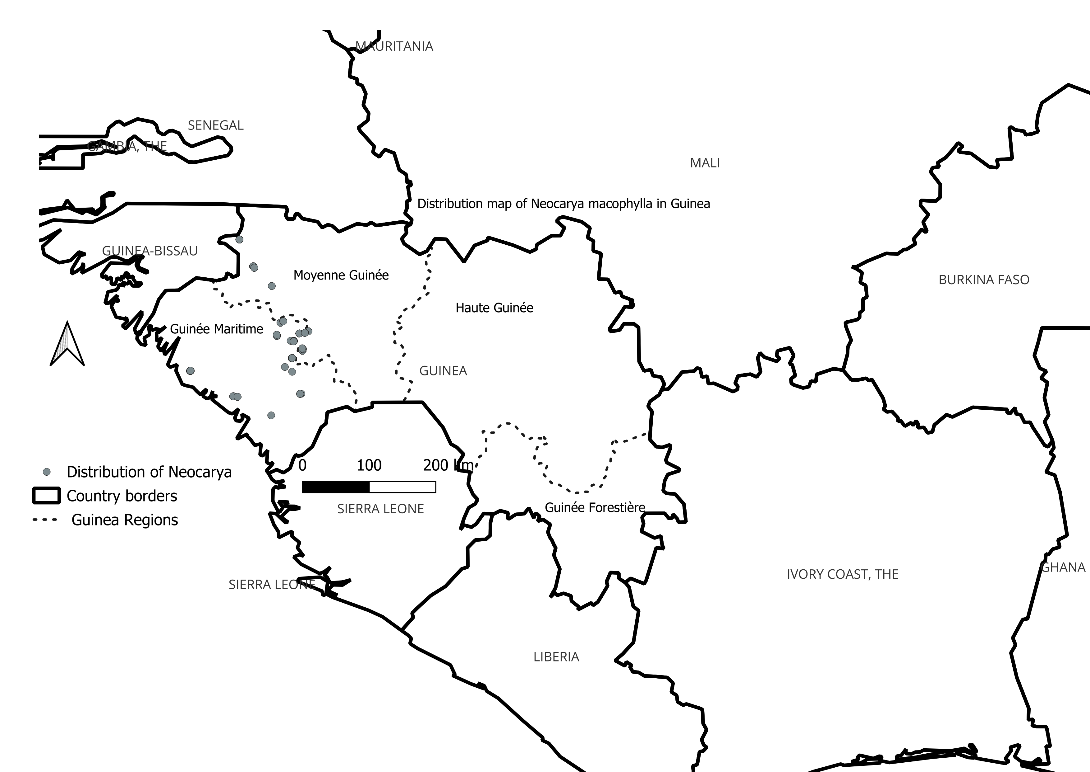
Based on field observations during recent surveys in Guinea, we found that the size of the sub-populations of *Neocarya macrophylla* at sites in Guinea varies from 3 to 150 trees. The greatest number of individuals per site were recorded west of the Fouta, with 100 or more trees per site in Pita and Gaoual Prefectures.

**Uses:** *Neocarya macrophylla* is a versatile tree with many uses. Fruit and seed are edible and appreciated by local communities, who wild harvest quantities. Seed and especially fruit are often sold in local markets, and the seeds are available on sale much of the year-round. The seeds, bark, root, and leaves have various medicinal applications. The plant provides useful timber, and several other products.

Figure 1: Map of distribution in Guinea based on verified herbarium specimens.

*Neocarya macrophylla* fruit (in fact the endocarp) is opened with a machete to extract the two seeds after placing it on a piece of plywood, which is slow and laborious and with resulting risks of injury. As a result, the quantity of seeds produced in Guinea is smaller than it might be. This is similar to findings from Balla *et al*. (2008). In Niger, the endocarp fragments are also used as fuel.

A recent analysis showed that *Neocarya* *macrophylla* seeds have a higher protein content than peanut (groundnut) per weight, which could be a good source of nutrition for local communities (Howes, unpublished data). Physicochemical studies carried out in Niger and Guinea confirm the presence of antioxidants in oils extracted from *Neocarya macrophylla*. (Diaby *et al*. 2016).

In Guinea, the roots, leaves, bark, and oil of this tree are widely used in traditional pharmacopoeia. They are used to prepare a decoction for stomach aches, dysentery, blood pressure and haemorrhoids. The seeds or kernels are either eaten raw, crushed, and used in sauces as condiments, or used for traditional oil extraction (United Purpose 2023).

The fruits and the seeds are eaten fresh. The seeds (in Nigeria) are boiled with cereal (corn or cassava flour) (Yusuf *et al.* 2024). *Neocarya macrophylla* seeds are roasted and eaten like cashews or almonds. *Neocarya macrophylla* is traditionally used as a food for medicinal, spiritual, and industrial purposes in Nigeria (Yusuf *et al.* 2024). It is also used as soap, dye, glue, fodder, termite repellent, firewood, and construction material. Several studies on the physicochemical, nutritional, phytochemical, and pharmacological content have validated the benefits of *N. macrophylla* for humans as a food, cosmetic and pharmaceutical. The main bioactive constituents identified to date in the plant are steroids and flavonoids (Diaby *et al*. 2016).

In the northern regions of Nigeria, the species is widely used to treat many illnesses such as asthma, skin infections, wounds, dysentery, inflammations, lung disorders, ear, and eye infections, (Yusuf *et al.* 2024). In traditional Nigerian medicine, the fruit is used to treat diarrhoea, while the seeds are used to heal wounds. (Yusuf *et al.* 2024).

Stem bark is used to treat conjunctivitis, pain, tooth decay, respiratory disorders, cancer and snakebite (Yusuf *et al.* 2024). Decoctions of leaves and bark are used as mouthwash, against internal disorders and for inflamed eyes (Yusuf *et al.* 2024).

In Senegal, the branches of the species are burned, and the smoke inhaled as a remedy for snakebites (Audu *et al*. 2005). The roots are used as antivenom, haemostatic agents and in the treatment of circumcision and wounds (Audu *et al*. 2005).

The fruit presents no known risks, meaning it is safe for human consumption.

### Identification of threats

This species is a savannah tree of sandy soils. The main threat for this species in the Maritime and Middle Guinea regions is uncontrolled bushfires, sand quarrying, logging, and the use of its bark in traditional medicine. Pressure on *Neocarya macrophylla* habitat is high. The bowé and plains are the zones most affected by bush fires in both regions. Cattle herders looking for new shoots to feed their domestic animals (cattle) set fire to the savannahs, wreaking havoc on all the species found there. Urbanization is another factor threatening *N. macrophylla*.

In Niger, in the Dallol Bosso region, analyses showed that excessive logging and unfavourable climatic conditions are the main potential threat factors for *Neocarya macrophylla*. Observation of the threats reveals that the greatest threat to *N. macrophylla* in Niger is because the women who produce firewood obtain it by improperly cutting the roots of living plants, which they then dry to boil water for making natron. Erosion contributes to the uprooting of trees (Aboubacar et al. 2018).

### Species management and conservation strategies:

*In situ* protection: No conservation actions are in place for this species in Guinea. Village communities protect this species in some areas because its fruit is a source of income for them. No *Neocarya macrophylla* site is protected by a state-recognized conservation act, nor by nature conservationists in Guinea. One collection has been reported in the classified Forest of Pic de Fon, a zone of iron mineralization, however, it is likely that this is a misidentification. Only the Mount Gangan and Tonkoyah Tropical Important Plant Areas (TIPAs) contain this species, but they are not formally protected (project run by RBG Kew and Herbier National de Guinea, Couch et al. 2019) (<http://www.herbierguinee.org/conservation-des-arbres-menacees.html>).

Monitoring the TIPAs where the species occurs to support the survival of the species' is recommended, in addition to maintaining existing natural stands.

This species is found in national parks and reserves in other West African countries such as: Senegal, Serra Leone, Liberia, and Nigeria (Yusuf *et al.* 2024).

*Ex situ* protection: Two seed collections from Mali are held at the Millennium Seed Bank Partnership in the UK. Germination trials of stored seeds have not yet been attempted. Fruits of this species were collected during our field surveys in 2020. Some experiments have been carried out with fruits and seeds. Intact fruits produce seedlings more easily than do extracted seeds. An experiment to distribute seedlings was carried out at Kakiwondi forest in Guinea to promote the cultivation of *Necarya* in a locality outside its natural range. Monitoring is ongoing, but it is not looking a very successful, probably as this area has a higher rainfall than in the natural range of the species. It is recommended to experiment with enrichment planting at such as Mont Gangan where the species occurs naturally and which are in the proposed National Parks network.

Ex situ conservation is likely not needed at this stage as the number of wild individuals is likely many hundreds or thousands in Guinea, and the species is ranged over West Africa. Moreover, cultivation efforts outside the local range of the species in Guinea have been problematic in our experience.

Objectives and targets:

Objective 1: Understand the threats to the in-situ population and how to control them.

Analysis of a field study in Guinea in 2020 showed that natural regeneration at sites for the species throughout the country is very low, with juveniles (< 1 m tall) at <10% the number of mature trees, and at 7 of 17 sites studied, no juveniles present. Fire exclusion experimental plots are advised in stands of *Neocarya* to test the hypothesis that fire, believed to be the main threat (55% of interviewees) to the species, is indeed reducing natural regeneration, and not another factor.

If fire is confirmed as a major threat, then installation of fire breaks around natural stands could be considered to protect the species. Monitoring of a random selection of sites is advisable to determine if the population is stable or not in Guinea.

Objective 2: Raising awareness among the local communities of the importance of protecting this species.

Local communities to be made aware of the importance of this species and the potential benefits it can bring in connection to food security and resilience to climate change. This species is adapted to the Guinean climate and could provide a valuable food resource in place of introduced species such as cashew nuts (*Anacardium occidentale*).

This species could be included in replanting efforts in suitable areas across the region.

Legislation:

The species has been assessed against IUCN Red List criteria as NT (Near Threatened, Molmou in press). It is advisable that it is added to the Monographie Nationale de la Guinée and also the species appendix of the Guinea Forest Act.
